## Supplementary Figures for "Crystal Structure of the 4-Hydroxybutyryl-CoA Synthetase (ADP-forming) from Nitrosopumilus maritimus"

### Supplementary Fig. 1- A reduced version of the sequence alignment of Nmar_0206 and others within the superfamily.

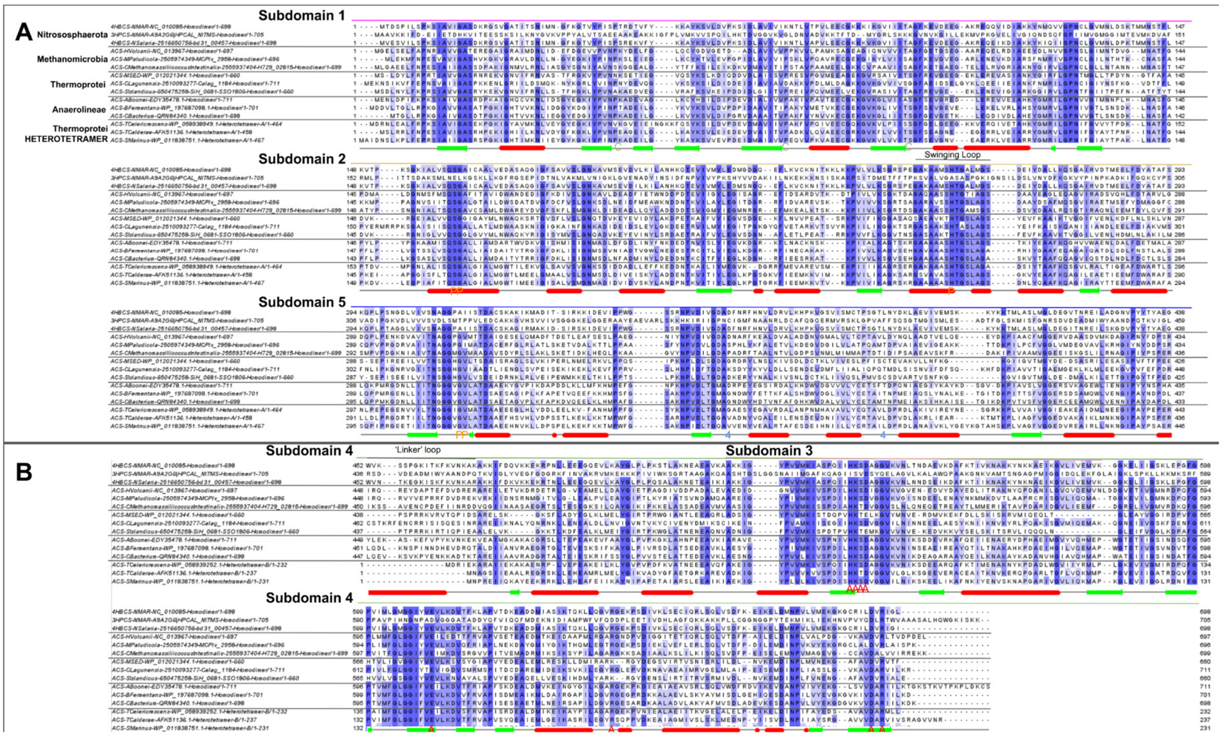


4 classes are present within the alignment including Nitrososphaeria, Methanomicrobia, Anaerolineae, and Thermoprotei. The Thermoprotei class displayed two sets of Acyl-CoA synthetases, one homodimeric structure like Nmar_0206 and the others, heteromeric like the *Ca.* Korarchaeum structure (Figs. 7 and 8). All sequences are available as supplementary files. Labels C, 4, P and A indicate possible Coenzyme, 4HB, Phosphate (Fig. 4) and ATP binding residues (Supplementary Fig. 3), respectively. The linker loop (Fig. 5) and swinging loop have additionally been labeled. A full list of sequences can be found in Supplementary Table 1.

### Supplementary Fig. 2- 4HB-CoA placed into binding site.

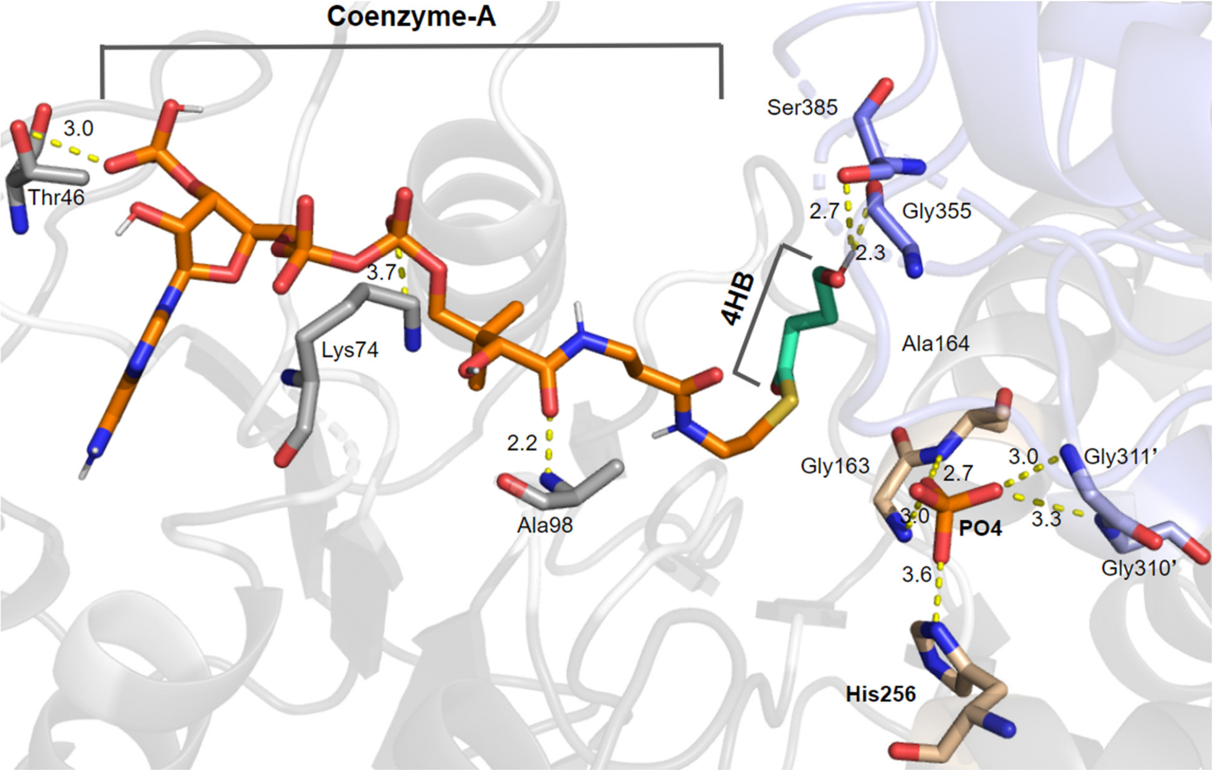


Using the 4YAK structure as a guide, 4HB-CoA was placed into its likely active site. Residues which could interact with the product are highlighted. The 4HB tail was manipulated looking for possible interactions for which Ser385 and Gly355 seemed possible candidates.

**Supplementary Fig. 3- ATP binding residues within Subdomain 3, the ATP-binding domain of Nmar_0206.**

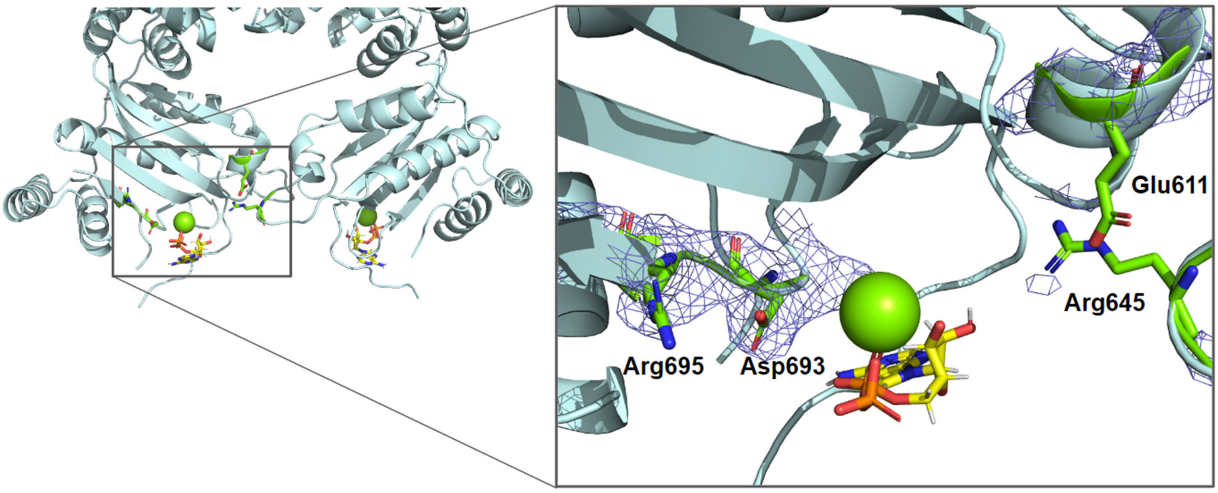


Although subdomain 4 is not present within our structure, residues Arg695 and Asp693 predicted by Weiße et. al (2016; corresponding Arg226 and Asp224) to interact with ATP are present. ADPCPhere (yellow) was copied from a superimposed pdb: 4XYM for rough estimation.

### Supplementary Fig. 4- Structural representations of the ACD and Nmar_0206 reaction mechanism.

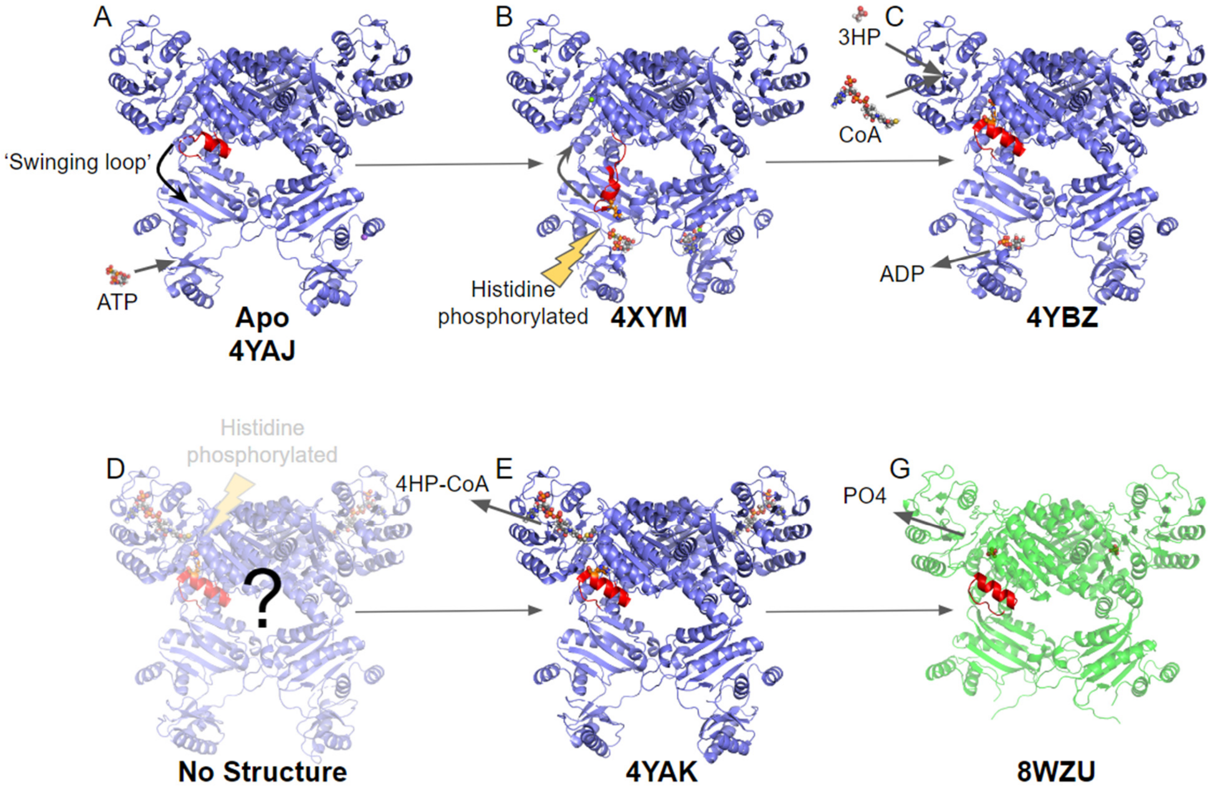


Blue structures indicate those produced by Weiße et al. while green is this manuscript's Nmar_0206 structure. The swinging loop is indicated in red. A) Shows the conformational state of the unbound apo-enzyme. B) Upon binding the ATP, the swinging loop moves the conserved His256 nearby the ATP-grasping domain. C) The swinging loop returns to the active site following the phosphorylation of His256 which may support the binding of further substrates. D) A structural intermediate showing the transient Acyl-phosphohistidine formation has yet to be determined. E) A structure with the bound Acyl-CoA product just prior to release. G) As phosphate is further within a pocket where the acetylation event takes place, the Nmar_0206 structure may represent a state just prior to the release of a final substrate.
