## Supplementary Table for "Crystal Structure of the 4-Hydroxybutyryl-CoA Synthetase (ADP-forming) from Nitrosopumilus maritimus"

### Supplementary Table 1: Sequence of Nmar_0206 homologues.

| Organism Name | Protein name | Accession Number | Domain/Phylum/Class |
|---|---|---|---|
| *Aciduliprofundum boonei* | acetyl coenzyme A synthetase (ADP forming) | WP_008084432.1 | Archaea/Euryarchaeota/DHVE2 |
| *Bellilinea caldifistulae* | acetyl-CoA synthetase | WP_061917260.1 | Bacteria/Chloroflexota/Anaerolineae |
| *Brevefilum fermentans* | acetate–CoA ligase family protein | WP_197687098.1 | Bacteria/Chloroflexota/Anaerolineae |
| *Candidatus Eisenbacteria* | acetate–CoA ligase family protein | WP_008084432.1 | Bacteria/Chloroflexota/Anaerolineae |
| *Chloroflexi bacterium* | acetate–CoA ligase family protein | NLG99206.1 | Bacteria/Chloroflexota |
| *Leptolinea sp.* | acetate–CoA ligase family protein | NMB56267.1 | Bacteria/Chloroflexota/Anaerolineae |
| *Leptolinea tardivitalis* | acetate–CoA ligase family protein | WP_062421259.1 | Bacteria/Chloroflexota/Anaerolineae |
| *Levilinea saccharolytica* | acetate–CoA ligase family protein | WP_062418633.1 | Bacteria/Chloroflexota/Anaerolineae |
| *Candidatus Methanomassiliicoccus intestinalis* | acetyltransferase | WP_020448504.1 | Archaea/Euryarchaeota/Thermoplasmata |
| *Haloferax volcanii* | acetyl-CoA synthetase | WP_004043943.1 | Archaea/Euryarchaeota/Halobacteria |
| *Methanobacterium congolense* | acetate–CoA ligase family protein | WP_071906649.1 | Archaea/Euryarchaeota/Methanobacteria |
| *Methanobacterium formicicum* | acetate–CoA ligase family protein | WP_004030442.1 | Archaea/Euryarchaeota/Methanobacteria |
| *Methanobacterium petrolearium* | acetate–CoA ligase family protein | WP_209625374.1 | Archaea/Euryarchaeota/Methanobacteria |
| *Methanobacterium sp.* | CoA-binding domain-containing protein | WP_292756096.1 | Archaea/Euryarchaeota/Methanobacteria |
| *Methanobacterium subterraneum* | acetate–CoA ligase family protein | WP_169032909.1 | Archaea/Euryarchaeota/Methanobacteria |
| *Methanocella arvoryzae* | acetate-CoA ligase | WP_ 012036318.1 | Archaea/Euryarchaeota/Methanomicrobia |

| | | | |
|---|---|---|---|
| *Methanocella conradii* | acetate-CoA ligase | WP_014404925.1 | Archaea/Euryarchaeota/Methanomicrobia |
| *Methanocella paludicola* | acetate–CoA ligase family protein | WP_012901660.1 | Archaea/Euryarchaeota/Methanomicrobia |
| *Methanococcoides burtonii* | acetate-CoA ligase | WP_011499410.1 | Archaea/Euryarchaeota/Methanomicrobia |
| *Methanococcoides methylutens* | acetate-CoA ligase | WP_048205925.1 | Archaea/Euryarchaeota/Methanomicrobia |
| *Methanococcoides orientis* | acetate–CoA ligase family protein | WP_233084021.1 | Archaea/Euryarchaeota/Methanomicrobia |
| *Methanococcoides seepicolus* | acetate–CoA ligase family protein | WP_250868540.1 | Archaea/Euryarchaeota/Methanomicrobia |
| *Methanohalobium evestigatum* | acetyl coenzyme A synthetase | WP_013195317.1 | Archaea/Euryarchaeota/Methanomicrobia |
| *Methanohalophilus euhalobius* | acetate-CoA ligase | WP_096712683.1 | Archaea/Euryarchaeota/Methanomicrobia |
| *Methanohalophilus levihalophilus* | acetate–CoA ligase family protein | WP_209682318.1 | Archaea/Euryarchaeota/Methanomicrobia |
| *Methanohalophilus mahii* | acetate-CoA ligase | WP_013038092.1 | Archaea/Euryarchaeota/Methanomicrobia |
| *Methanohalophilus portucalensis* | acetate-CoA ligase | WP_072360490.1 | Archaea/Euryarchaeota/Methanomicrobia |
| *Methanohalophilus profundi* | acetate-CoA ligase | WP_129598452.1 | Archaea/Euryarchaeota/Methanomicrobia |
| *Methanolobus bombayensis* | acetate–CoA ligase family protein | WP_209618454.1 | Archaea/Euryarchaeota/Methanomicrobia |
| *Methanolobus chelungpuianus* | acetate–CoA ligase family protein | WP_256621754.1 | Archaea/Euryarchaeota/Methanomicrobia |
| *Methanolobus halotolerans* | acetate-CoA ligase | WP_135389970.1 | Archaea/Euryarchaeota/Methanomicrobia |
| *Methanolobus profundi* | acetate-CoA ligase | WP_091937667.1 | Archaea/Euryarchaeota/Methanomicrobia |
| *Methanolobus psychrophilus* | acetyl coenzyme A synthetase | AFV23390.1 | Archaea/Euryarchaeota/Methanomicrobia |
| *Methanolobus psychrotolerans* | acetate-CoA ligase | WP_094228889.1 | Archaea/Euryarchaeota/Methanomicrobia |
| *Methanolobus tindarius* | acetate-CoA ligase | WP_023845745.1 | Archaea/Euryarchaeota/Metha |

| | | | |
|---|---|---|---|
| | | | nomicrobia |
| *Methanolobus zinderi* | acetate-CoA ligase | WP 176964306.1 | Archaea/Euryarchaeota/Methanomicrobia |
| *Methanomethylovorans hollandica* | acetate–CoA ligase family protein | WP_015324018.1 | Archaea/Euryarchaeota/Methanomicrobia |
| *Methanosalsum natronophilum* | acetate–CoA ligase family protein | WP_259134952.1 | Archaea/Euryarchaeota/Methanomicrobia |
| *Methanosalsum zhilinae* | acetate-CoA ligase | WP_013898132.1 | Archaea/Euryarchaeota/Methanomicrobia |
| *Armatimonadetes bacterium* | acetate–CoA ligase family protein | MBC8157135.1 | Bacteria/Armatimonadota/undefined |
| *Candidatus Nitrosarchaeum limnium* | acetate–CoA ligase family protein | WP_010190675.1 | Archaea/Nitrososphaerota/Nitrososphaeria |
| *Candidatus Nitrosopelagicus brevis* | CoA-binding protein | NMI83880.1 | Archaea/Nitrososphaerota/Nitrososphaeria |
| *Candidatus Nitrosopelagicus sp.* | acylCoA synthetase | GIT55543.1 | Archaea/Nitrososphaerota/Nitrososphaeria |
| *Candidatus Nitrosopolaris wilkensis* | acylCoA synthetase | PWU81816.1 | Archaea/Nitrososphaerota/Nitrososphaeria |
| *Candidatus Nitrosopumilus koreensis* | 3-hydroxypropionyl-CoA synthetase (ADP-forming) | WP_007551149.1 | Archaea/Nitrososphaerota/Nitrososphaeria |
| *Candidatus Nitrosopumilus sediminis* | acetate-CoA ligase | WP_014964350.1 | Archaea/Nitrososphaerota/Nitrososphaeria |
| *Candidatus Nitrososphaera evergladensis* | 4-hydroxybutyryl-CoA synthetase (ADP-forming) | WP_148699949.1 | Archaea/Nitrososphaerota/Nitrososphaeria |
| *Candidatus Nitrososphaera gargensis* | acetate–CoA ligase family protein | WP_015020118.1 | Archaea/Nitrososphaerota/Nitrososphaeria |
| *Candidatus Nitrosotalea devanaterra* | 4-hydroxybutyryl-CoA synthetase (ADP-forming) | CUR50945.1 | Archaea/Nitrososphaerota/Nitrososphaeria |
| *Candidatus Nitrosotalea bavarica* | acetate–CoA ligase family protein | WP_101477917.1 | Archaea/Nitrososphaerota/Nitrososphaeria |
| *Candidatus Nitrosotalea devanaterra* | 4-hydroxybutyryl-CoA synthetase (ADP-forming) | CUR50945.1 | Archaea/Nitrososphaerota/Nitrososphaeria |

| | | | |
|---|---|---|---|
| *Candidatus Nitrosotalea okcheonensis* | acetate–CoA ligase family protein | WP_157928055.1 | Archaea/Nitrososphaerota/Nitrososphaeria |
| *Candidatus Nitrosotalea sinensis* | acetate–CoA ligase family protein | WP_101008906.1 | Archaea/Nitrososphaerota/Nitrososphaeria |
| *Candidatus Nitrosotenuis aquarius* | acetate-CoA ligase | WP_100182658.1 | Archaea/Nitrososphaerota/Nitrososphaeria |
| *Candidatus Nitrosotenuis chungbukensis* | acetate–CoA ligase family protein | WP_042683670.1 | Archaea/Nitrososphaerota/Nitrososphaeria |
| *Candidatus Nitrosotenuis cloacae* | acetate–CoA ligase | WP_048186944.1 | Archaea/Nitrososphaerota/Nitrososphaeria |
| *Candidatus Nitrosotenuis sp.* | CoA-binding protein | TBR22739.1 | Archaea/Nitrososphaerota/Nitrososphaeria |
| *Candidatus Nitrosotenuis uzonensis* | acetate–CoA ligase family protein | WP_048194424.1 | Archaea/Nitrososphaerota/Nitrososphaeria |
| *Cenarchaeum symbiosum* | acyl-CoA synthetase (NDP- forming) | ABK76672.1 | Archaea/Nitrososphaerota/Nitrososphaeria |
| *Nitrosarchaeum koreense* | acetate–CoA ligase family protein | WP_007549553.1 | Archaea/Nitrososphaerota/Nitrososphaeria |
| *Nitrosopumilus cobalaminigenes* | acetate–CoA ligase | WP_179361083.1 | Archaea/Nitrososphaerota/Nitrososphaeria |
| *Nitrosopumilus maritimus (strain SCM1)* | 3-hydroxypropionyl-CoA synthetase (ADP-forming) | WP_012215692.1 | Archaea/Nitrososphaerota/Nitrososphaeria |
| *Nitrosopumilus maritimus (strain SCM1)* | 4-hydroxybutyryl-CoA synthetase (ADP-forming) | WP_012214589.1 | Archaea/Nitrososphaerota/Nitrososphaeria |
| *Nitrosopumilus oxyclinae* | acetate–CoA ligase family protein | WP_179362924.1 | Archaea/Nitrososphaerota/Nitrososphaeria |
| *Nitrosopumilus salaria* | 4-hydroxybutyryl-CoA synthetase (ADP-forming) | WP_008297726.1 | Archaea/Nitrososphaerota/Nitrososphaeria |
| *Nitrosopumilus sp.* | 4-hydroxybutyryl-CoA synthetase (ADP-forming) | RMW38333.1 | Archaea/Nitrososphaerota/Nitrososphaeria |
| *Nitrososphaera viennensis* | 4-hydroxybutyryl-CoA synthetase (ADP-forming) | WP_075055435.1 | Archaea/Nitrososphaerota/Nitrososphaeria |
| *Candidatus Nitrosotalea okcheonensis* | acetate–CoA ligase family protein | WP_157928055.1 | Archaea/Nitrososphaerota/Nitrososphaeria |

| | | | |
|---|---|---|---|
| *Candidatus Nitrosocosmicus oleophilus* | acetate–CoA ligase family protein | WP_196817532.1 | Archaea/Nitrososphaerota/Nitrososphaeria |
| *Staphylothermus hellenicus* | acetyl coenzyme A synthetase (ADP forming) | ADI31488.1 | Archaea/Thermoproteota/Thermoprotei |
| *Ignisphaera aggregans* | acetyl coenzyme A synthetase (ADP forming) | ADM28387.1 | Archaea/Thermoproteota/Thermoprotei |
| *Pyrolobus fumarii* | CoA-binding domain protein | AEM39269.1 | Archaea/Thermoproteota/Thermoprotei |
| *Thermogladius calderae* | CoA-binding domain protein | AFK51136.1 | Archaea/Thermoproteota/Thermoprotei |
| *Desulfurococcales archaeon* | CoA-binding protein | MCD6324124.1 | Archaea/Thermoproteota/Thermoprotei |
| *Candidatus Bathyarchaeota archaeon* | acetyl coenzyme A synthetase (ADP forming) | HDM23685.1 | Archaea/Bathyarchaeota/undefined |
| *Acidilobales archaeon* | acetyl coenzyme A synthetase (ADP forming) | HDN75497.1 | Archaea/Thermoproteota/Thermoprotei |
| *Fervidicoccaceae archaeon* | CoA-binding protein | MCC6010103.1 | Archaea/Thermoproteota/Thermoprotei |
| *Staphylothermus sp.* | CoA-binding protein | MCD6196563.1 | Archaea/Thermoproteota/Thermoprotei |
| *Desulfurococcales archaeon* | CoA-binding protein | MCD6488109.1 | Archaea/Thermoproteota/Thermoprotei |
| *Thermoprotei archaeon* | CoA-binding protein | MCI4396430.1 | Archaea/Thermoproteota/Thermoprotei |
| *Ignisphaera sp.* | CoA-binding protein | MCI4436598.1 | Archaea/Thermoproteota/Thermoprotei |
| *Crenarchaeota archaeon* | acetyl-CoA synthetase (ADP forming) | NPA97047.1 | Archaea/Thermoproteota/Thermoprotei |
| *Aigarchaeota archaeon NZ13 MG1* | acetyl-CoA synthetase (ADP forming) | PUA31671.1 | Archaea/Aigarchaeota/undefined |
| *Zestosphaera tikiterensis* | acetyl-CoA synthetase (ADP forming) | PUA33430.1 | Archaea/Thermoproteota/Thermoprotei |
| *Deltaproteobacteria bacterium* | acetyl-CoA synthetase (ADP forming) | RLB79341.1 | Archaea/Deltaproteobacteria/undefined |

| | | | |
|---|---|---|---|
| *Candidatus Verstraetearchaeota archaeon* | acetyl-CoA synthetase (ADP forming) | RLE53850.1 | Archaea/Verstraetearchaeota/undefined |
| *Thermoprotei archaeon* | acetyl-CoA synthetase (ADP forming) | RLE57312.1 | Archaea/Thermoproteota/Thermoprotei |
| *Hyperthermus butylicus* | CoA_binding protein | WP_011822483.1 | Archaea/Thermoproteota/Thermoprotei |
| *Staphylothermus marinus* | CoA_binding protein | WP 011838751.1 | Archaea/Thermoproteota/Thermoprotei |
| *Candidatus Korarchaeum cryptofilum* | CoA_binding protein | WP_012308855.1 | Archaea/Korarchaeota/Korarchaei |
| *Fervidicoccus fontis* | CoA_binding protein | WP_014557871.1 | Archaea/Thermoproteota/Thermoprotei |
| *Thermococcus cleftensis* | acetate-CoA ligase | WP_014789415.1 | Archaea/Thermoproteota/Thermoprotei |
| *Staphylothermus hellenicus* | CoA_binding protein | WP_052833590.1 | Archaea/Thermoproteota/Thermoprotei |
| *Thermofilum adornatum* | CoA_binding protein | WP_052886942.1 | Archaea/Thermoproteota/Thermoprotei |
| *Thermococcus celericrescens* | acetate-CoA ligase | WP_058938949.1 | Archaea/Thermoproteota/Thermoprotei |
| *Thermosphaera aggregans* | CoA_binding protein | WP_193435632.1 | Archaea/Thermoproteota/Thermoprotei |
| *Thermogladius calderae* | CoA_binding protein | WP_202945702.1 | Archaea/Thermoproteota/Thermoprotei |
| *Caldisphaera lagunensis* | acetyltransferase | WP_015232801.1 | Archaea/Thermoproteota/Thermoprotei |
| *Metallosphaera sedula* | acetyl-CoA synthetase (ADP forming) alpha domain containing protein | WP_240252923.1 | Archaea/Thermoproteota/Thermoprotei |
| *Sulfolobus islandicus* | acetyl-CoA synthetase (ADP forming) alpha domain containing protein | WP_012715981.1 | Archaea/Thermoproteota/Thermoprotei |
| *Sulfolobus solfataricus* | acetyl-CoA synthetase (ADP forming) alpha domain containing protein | WP_009992015.1 | Archaea/Thermoproteota/Thermoprotei |

| | | | |
|---|---|---|---|
| *Infirmifilum uzonense* | acetyltransferase | WP_191118496.1 | Archaea/Thermoproteota/Thermoprotei |
